# Fission yeast CK1 promotes DNA double-strand break repair through both homologous recombination and non-homologous end joining

**DOI:** 10.1101/2023.04.27.538600

**Authors:** Sierra N. Cullati, Eric Zhang, Yufan Shan, Rodrigo X. Guillen, Jun-Song Chen, Jose Navarrete-Perea, Zachary C. Elmore, Liping Ren, Steven P. Gygi, Kathleen L. Gould

**Affiliations:** Department of Cell and Developmental Biology, Vanderbilt University School of Medicine, Nashville, TN, USA; Department of Cell Biology, Harvard Medical School, Boston, MA, USA; Columbia University Medical Center, New York, NY, USA; Department of Surgery, Duke University School of Medicine, Durham, NC, USA

**Keywords:** CK1, casein kinase 1, *Schizosaccharomyces pombe*, DNA repair, non-homologous end joining, homologous recombination, INO80, Arp8, phosphoproteomics

## Abstract

The CK1 family are conserved serine/threonine kinases with numerous substrates and cellular functions. The fission yeast CK1 orthologues Hhp1 and Hhp2 were first characterized as regulators of DNA repair, but the mechanism(s) by which CK1 activity promotes DNA repair had not been investigated. Here, we found that deleting Hhp1 and Hhp2 or inhibiting CK1 catalytic activities in yeast or in human cells activated the DNA damage checkpoint due to persistent double-strand breaks (DSBs). The primary pathways to repair DSBs, homologous recombination and non-homologous end joining, were both less efficient in cells lacking Hhp1 and Hhp2 activity. In order to understand how Hhp1 and Hhp2 promote DSB repair, we identified new substrates using quantitative phosphoproteomics. We confirmed that Arp8, a component of the INO80 chromatin remodeling complex, is a bona fide substrate of Hhp1 and Hhp2 that is important for DSB repair. Our data suggest that Hhp1 and Hhp2 facilitate DSB repair by phosphorylating multiple substrates, including Arp8.

## Introduction

DNA damage poses a constant threat to cells by introducing alterations to the genetic code, impeding DNA replication, and contributing to genomic instability. In addition to external hazards such as UV light and genotoxic chemicals, cells frequently experience DNA damage during normal cellular processes such as replication, transcription, and meiosis. Uncontrolled DNA damage is detrimental to cell survival, and in the case of metazoans, can contribute to cancer progression and therapeutic resistance. A complex arsenal of DNA repair machinery and signaling proteins make up the DNA damage response (DDR), which is coupled to cell cycle checkpoints to allow cells to recognize DNA damage, stall the cell cycle, and repair that damage prior to replication or cell division. DNA double-stranded breaks (DSBs) are a particularly cytotoxic type of DNA damage, capable of not only obstructing DNA replication and gene expression, but also causing the loss of genetic information or instigating gross chromosomal rearrangements (Cannan and Pederson, 2016). The two primary DSB repair pathways are homologous recombination (HR), which requires a second copy of DNA for homology-directed repair, and non-homologous end joining (NHEJ), which can directly ligate broken DNA ends, in addition to alternative repair pathways such as microhomology-mediated end-joining and single-strand annealing.

CK1 enzymes comprise a ubiquitous, conserved family of protein kinases with roles in many cellular processes, including the cell cycle, endocytosis, circadian rhythms, and stress responses (Knippschild et al., 2014). The mammalian CK1 family has seven isoforms (α1, α2, γ1, γ2, γ3, δ, ε) that are present to some degree in nearly every tissue type and cellular compartment, and they have hundreds of substrates, including some involved in the DDR such as p53, Mdm2, and claspin (Dumaz et al., 1999; Higashimoto et al., 2000; Inuzuka et al., 2010; Knippschild et al., 1997; Meng et al., 2011; Sakaguchi et al., 2000; Venerando et al., 2010; Winter et al., 2004). In *Schizosaccharomyces pombe*, Hhp1 and Hhp2 are orthologues of mammalian CK1δ and CK1ε, as well as *Saccharomyces cerevisiae* Hrr25 (Hoekstra et al., 1994). Hhp1 and Hhp2 are important for DSB repair (Dhillon and Hoekstra, 1994), and cells missing one or both genes are sensitive to an array of DNA damaging agents (Bimbó et al., 2005; Chen et al., 2015; Dhillon and Hoekstra, 1994).

Furthermore, deletion of *hhp1* alone or in combination with *hhp2* activates one or more cell cycle checkpoints. Fission yeast grow from the cell tips and divide medially at a set cell size; in this way, growth is coupled to the cell cycle, and cell cycle stage is closely correlated to cell length (Mitchison and Nurse, 1985). When a cell cycle checkpoint has been activated, growth continues during the G2 delay, resulting in cells that divide at a longer size than usual. *hhp1Δ* and *hhp1Δ hhp2Δ* cells have an increased length at septation indicating activation of a checkpoint (Dhillon and Hoekstra, 1994; Hayles et al., 2013). However, the mechanism(s) by which CK1 promotes DNA repair or activates this checkpoint are not understood.

In this study, we demonstrate that yeast and human cells lacking CK1 catalytic activity trigger the DNA damage checkpoint in response to persistent DSBs. We find that inhibiting Hhp1 and Hhp2 in *S. pombe* significantly impairs both HR and NHEJ, suggesting that these kinases phosphorylate multiple substrates to promote DSB repair through different pathways. We employed quantitative phosphoproteomics to identify Hhp1 and Hhp2 substrates contributing to DSB repair and validated one new substrate, Arp8, which is phosphorylated by Hhp1 and Hhp2 on four serines in its N-terminus.

Arp8 is a subunit of the INO80 chromatin remodeling complex, which evicts and slides nucleosomes around DSBs to promote end resection, an early step in DSB repair (Brahma et al., 2018; Gospodinov et al., 2011; Lademann et al., 2017; van Attikum et al., 2007). Because DNA repair pathways and CK1 enzymes are conserved between fission yeast and other eukaryotes, we predict that our findings will yield insight into how CK1 promotes DNA repair in human systems.

## Results

### Loss of Hhp1 and Hhp2 kinase activity triggers the DNA damage checkpoint

*hhp1Δ* and *hhp1Δ hhp2Δ* strains exhibit abnormally long cell lengths at septation and overall slower growth compared to *wildtype* (Dhillon and Hoekstra, 1994; Hayles et al., 2013). Deletion of *hhp2* alone does not result in these phenotypes (Dhillon and Hoekstra, 1994; Hayles et al., 2013), likely due to compensation by Hhp1, which is more abundant (Elmore et al., 2018). Increased length of *S. pombe* cells at septation results from a delay in mitotic entry (Mitchison and Nurse, 1985), and we reasoned that this may be due to activation of the DNA damage checkpoint because deletion of *hhp1* or both *hhp1* and *hhp2* impairs DSB repair (Dhillon and Hoekstra, 1994). Indeed, we observed that even without exposure to external DNA damaging agents *hhp1Δ* and *hhp1Δ hhp2Δ* cells had high levels of DSBs (Figure 1A-B) marked by the presence of Rad52-GFP. The HR protein Rad52 localizes diffusely in the nucleus under normal conditions, while in response to DSBs, Rad52-GFP forms bright repair foci (Alabert et al., 2009; Doe et al., 2004; Kim et al., 2000; Meister et al., 2005; van den Bosch M et al., 2001). We detected Rad52-GFP in nuclear foci across all stages of the cell cycle (Figure 1A), which is evidence of persistent DSBs. Rad52-GFP protein levels were unchanged in all strains examined (Supplemental Figure 1A).

**Figure 1:**
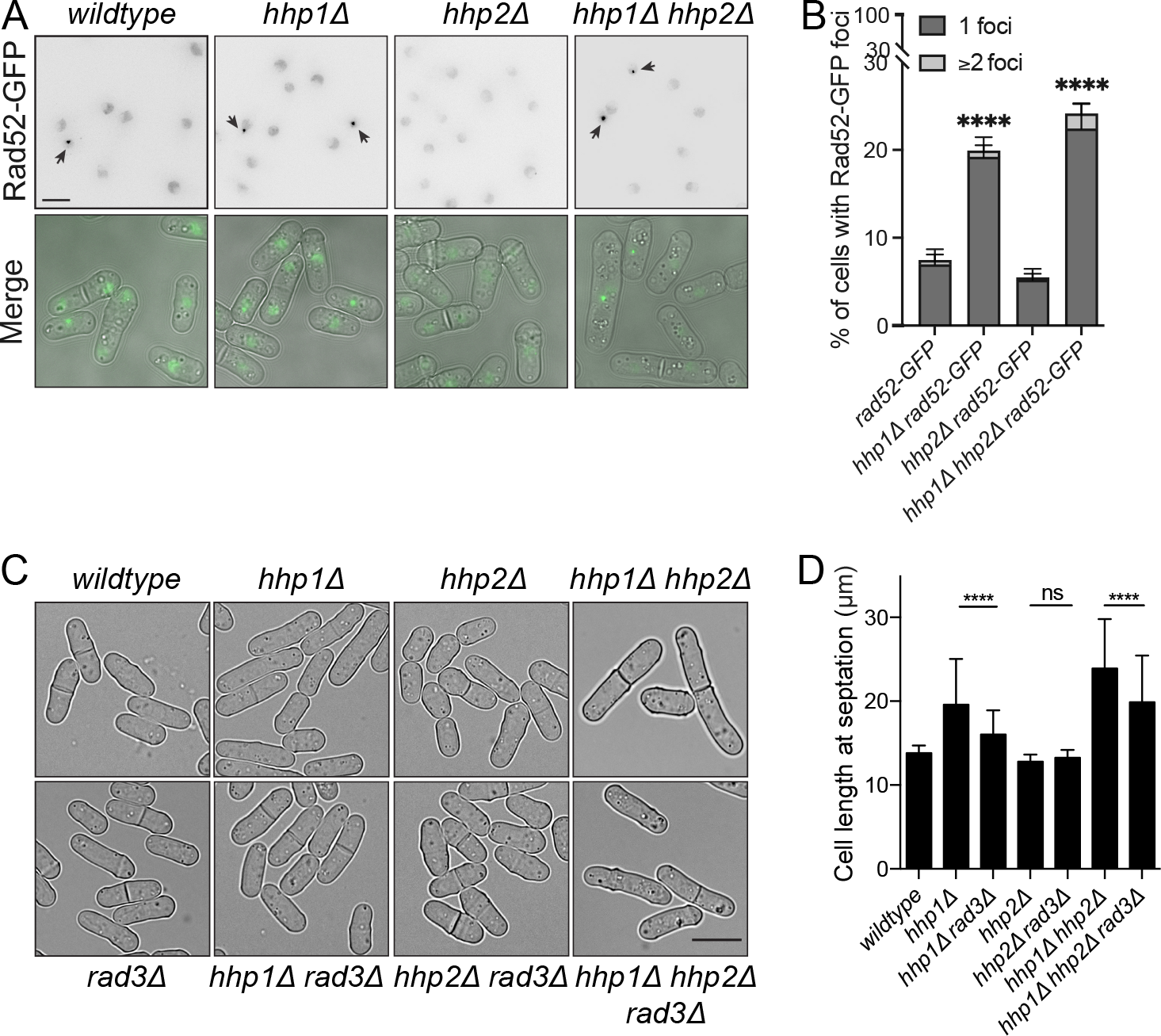
Deletion of *hhp1* alone or in combination with *hhp2* activates *rad3*- dependent DNA damage checkpoints. (A-B) *hhp1Δ* and *hhp1Δ hhp2Δ* strains exhibit increased basal levels of Rad52-GFP foci compared to *wildtype*. (A) Live-cell images of the indicated strains. Merge represents DIC and GFP channels. Rad52-GFP images are max projections. DIC images are single medial slices. (B) Rad52-GFP foci in > 600 cells for each strain were counted over 2 to 4 biological replicates. Error bars indicate ± 95% CI. ****, p < 0.0001 by Chi-square. (C) DIC live-cell images of the indicated strains. (D) Length at septation was quantified for cells imaged as in C. For *wildtype* and *hhp1Δ hhp2Δ* strains, n≥40 cells in each of 6 replicates. For all other strains, n ≥ 40 cells in each of 3 replicates. Bars represent means ± SD. ****, p < 0.0001 by one-way ANOVA; ns, not significant. Scale bars: 10 μm.

To test whether these persistent DSBs activated the DNA damage checkpoint leading to increased cell length, we deleted the master checkpoint kinase *rad3* in combination with *hhp1Δ, hhp2Δ,* or *hhp1Δ hhp2Δ*. Deleting *rad3* significantly reduced the length of *hhp1Δ* and *hhp1Δ hhp2Δ* cells (Figure 1C-D) but did not rescue growth (Supplemental Figure 1B). Further supporting the idea that activation of the DNA damage checkpoint delays mitotic entry in *hhp1Δ* and *hhp1Δ hhp2Δ* cells, deletion of the effector kinases Chk1 and Cds1, which are downstream of Rad3, also suppressed the elongated cell length phenotype but not cell growth (Supplemental Figure 1C-H).

Because eliminating the DNA damage checkpoint did not fully reduce *hhp1Δ hhp2Δ* cells to the *wildtype* length at septation, we considered whether another cell cycle checkpoint was also activated in these cells. In the presence of cellular stress, MAP kinase Sty1 activates Srk1 to inhibit mitotic entry (López-Avilés et al., 2005). We found that *hhp1Δ srk1Δ* and *hhp1Δ hhp2Δ srk1Δ* had cell lengths similar to those of *hhp1Δ* and *hhp1Δ hhp2Δ*, respectively (Supplemental Figure 2A-B). We were unable to construct strains deleting *srk1, rad3, hhp1*, and/or *hhp2*, so we utilized *hhp1-M84G* and *hhp2-M85G* alleles that are specifically inhibited by the ATP analogue 1NM-PP1 (denoted *hhp1-as* and *hhp2-as*) (Cullati et al., 2022; Gregan et al., 2007; Johnson et al., 2013). After 5 hours of treatment with 25 μM 1NM-PP1, *hhp1-as hhp2-as* cells were long compared to controls (Supplemental Figure 2C). The length of *hhp1-as hhp2-as* cells was reduced by deleting *rad3* but not *srk1* alone, and the deletion of both *rad3* and *srk1* was no different than the *rad3* deletion (Supplemental Figure 2C). Thus, we propose that DNA damage checkpoint activation is the primary cause of G2 delay in cells with impaired Hhp1 and Hhp2 function.

*hhp1Δ, hhp2Δ,* and *hhp1Δ hhp2Δ* cells are sensitive to a broad spectrum of exogenous DNA damage sources, including agents that modify bases (e.g. methane methylsulfonate, MMS), obstruct progression of the replication fork (e.g. hydroxyurea, HU), and induce DNA breaks (e.g. camptothecin, CPT) (Bimbó et al., 2005; Chen et al., 2015; Dhillon and Hoekstra, 1994) (Supplemental Figure 3A). We have found that this is specifically due to loss of Hhp1 and Hhp2 kinase activity, as kinase-dead mutants are also sensitive to these agents (Supplemental Figure 3B). Furthermore, *hhp1-as hhp2-as* cells grow normally on rich media (Supplemental Figure 3C), but when treated with 1NM-PP1, they grow slowly and exhibit similar DNA damage sensitivities (Figure 2A).

**Figure 2:**
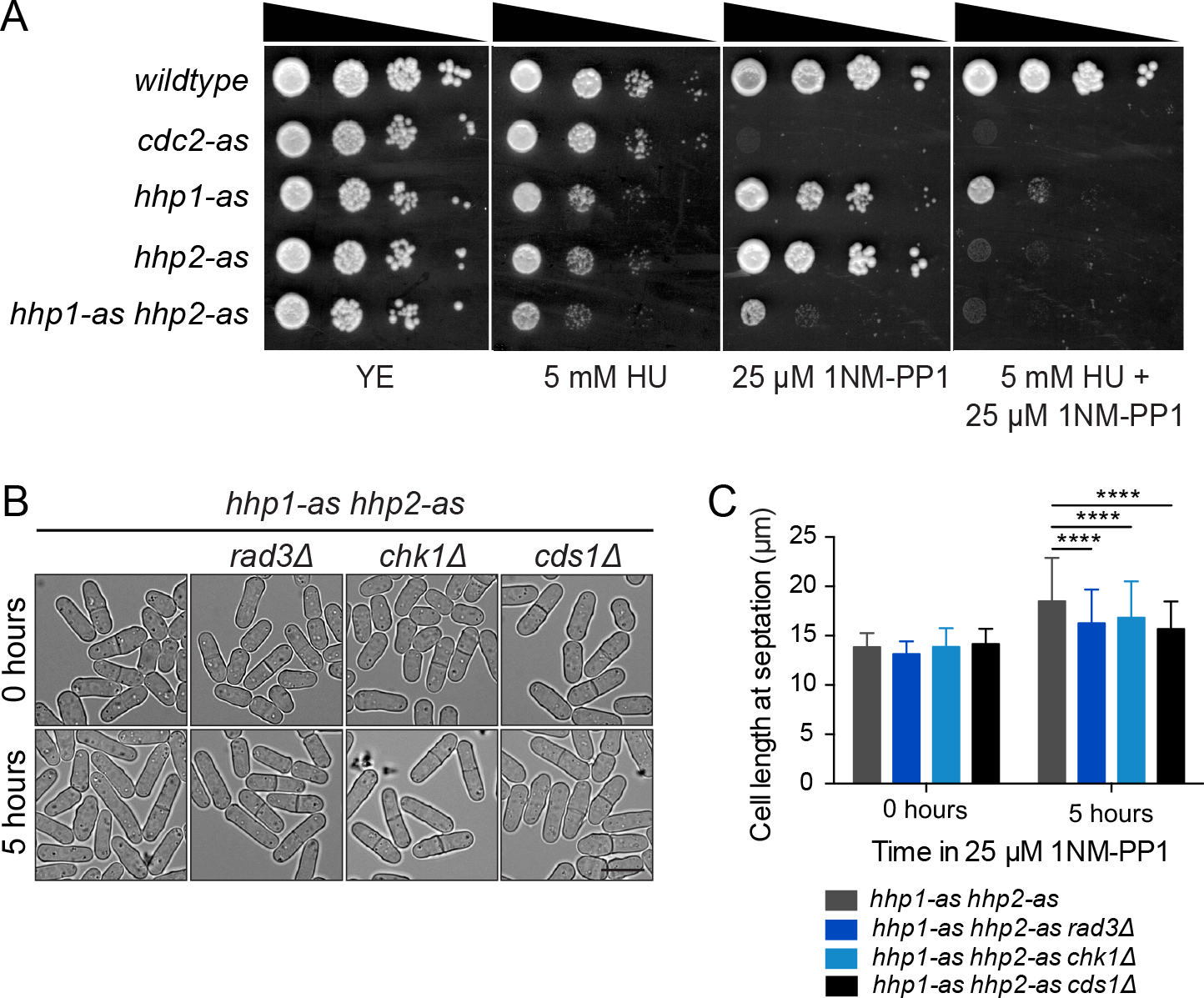
Loss of Hhp1 and Hhp2 activity triggers the DNA damage checkpoint. (A) Serial 10-fold dilutions of the indicated strains were spotted on YE with or without 1NM-PP1 and hydroxyurea (HU) and incubated at 32°C. *cdc2-as* was used as a positive control for inhibitor effectiveness; inhibition of Cdc2 is lethal. (B) DIC live-cell images of the indicated strains at 0 and 5 h after treatment with 25 μM 1NM-PP1. (C) Length at septation was quantified for cells imaged as in B, n ≥ 40 cells in each of 3 replicates. Bars represent means ± SD. ****, p < 0.0001 by one-way ANOVA; ns, not significant.

When we inhibited Hhp1 and Hhp2 activity in combination with deletion of DNA damage checkpoint kinases, we saw a similar suppression of the cell length phenotype as in the deletion mutants (Figure 2B-C). This suggests that like in *hhp1Δ* and *hhp1Δ hhp2Δ* cells, the DNA damage checkpoint is activated when *hhp1-as hhp2-as* kinase activity is inhibited for a shorter period of time.

### CK1 kinase activity promotes DSB repair in both yeast and human cells

The increase in DSBs and sensitivity to genotoxic stress that we observed in cells lacking Hhp1 and Hhp2 activity could be due to increased DNA damage and/or failure to repair the damage. To examine these possibilities, we used HU to both synchronize cells in S-phase and to induce replication stress. *hhp1-as hhp2-as rad52- GFP* cells were then released into media without HU but containing 25 μM 1NM-PP1 or DMSO as a control, and cells were imaged over time to measure the frequency of Rad52-GFP foci (Figure 3A-B). Rad52-GFP rapidly localized to foci following release from HU, and cells with or without 1NM-PP1 treatment exhibited the same frequency of foci (Figure 3B), suggesting that cells experience the same amount of DNA damage with and without Hhp1 and Hhp2 activity. However, while Rad52-GFP foci largely resolved within ∼2.5 hours in uninhibited cells, they were maintained for more than 5 hours in the presence of inhibitor (Figure 3A-B). These data indicate that cells lacking Hhp1 and Hhp2 kinase activity can still recognize DSBs and initiate a checkpoint response, but they are unable to complete DSB repair.

**Figure 3:**
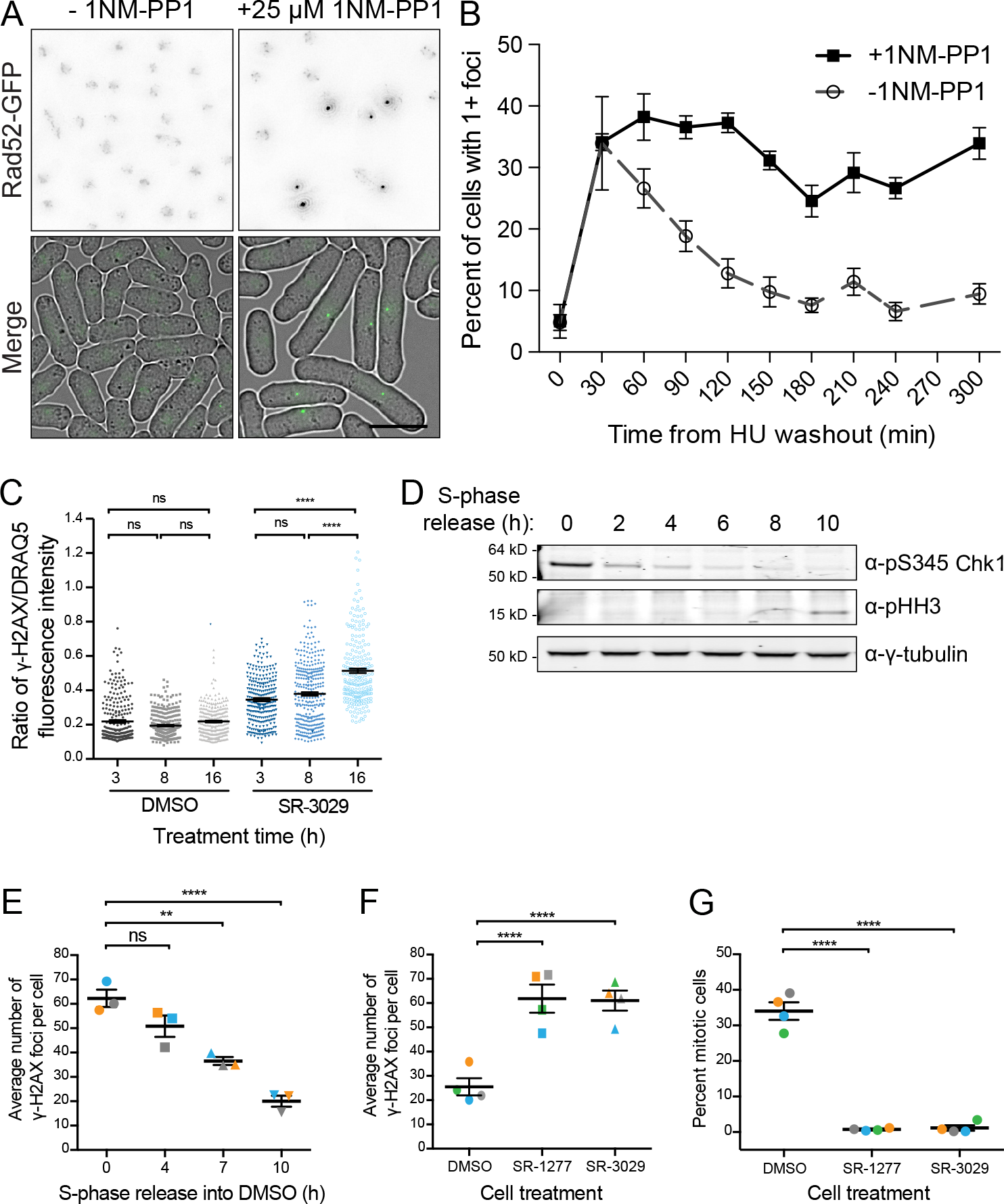
CK1 kinase activity promotes DSB repair in both yeast and human cells. (A-B) *hhp1-as hhp2-as rad52-GFP* cells were synchronized in 12 mM HU for 3 h then released into YE with or without 25 μM 1NM-PP1. Samples were fixed and imaged at the indicated timepoints. (A) Fluorescence microscopy and DIC images of the indicated strains after 240 min. Rad52-GFP images are max projections. DIC images are single medial slices. Scale bar: 10 μm. (B) Rad52-GFP foci were counted at the indicated timepoints using sum projections of cells imaged as in A. Data are means ± SEM from 3 replicates of n ≥ 150 each. (C) Asynchronous HeLa cells were treated with DMSO or 0.5 μM SR-3029 (CK1δ and CK1ε inhibitor) for the indicated times. Cells were fixed and stained for γH2AX as a marker of DSBs and DRAQ5 fluorescent probe for total DNA content. (D-G) HeLa cells were synchronized in S-phase using a thymidine-aphidicolin block. (D) Western blot of whole cell lysates with the indicated antibodies after release from S-phase arrest into DMSO. (E) Average number of γH2AX foci per cell after release from an S-phase arrest into DMSO. (F) Average number of γH2AX foci per cell 10 h after release from an S-phase arrest into DMSO, 0.5 μM SR-1227, or 0.5 μM SR- 3029. (G) Mitotic index 10 h after release from an S-phase arrest into DMSO, 0.5 μM SR-1227, or 0.5 μM SR-3029.

Because CK1 enzymes and the processes of DNA repair are conserved from yeast to human, we asked whether a similar DNA repair defect occurred when CK1δ and CK1ε, the human homologues of Hhp1 and Hhp2, were inhibited. γH2AX nuclear foci are a marker of DSBs in mammalian cells (Rogakou et al., 1998). We treated HeLa cells with SR-3029, a small molecule inhibitor of CK1δ and CK1ε (Bibian et al., 2013), and observed accumulation of γH2AX foci over time, even without any treatment to induce DNA damage (Figure 3C). This resembled our results in *hhp1Δ* and *hhp1Δ hhp2Δ* fission yeast (Figure 1A-B) and a previous report in *CSNK1D^-/-^* MEFs (Greer et al., 2017). Next we synchronized cells in S-phase and induced replication stress using a thymidine-aphidicolin block and release. As expected, cells initially activated the DDR (Figure 3D), as evidenced by activated Chk1 (Chen et al., 2000), due to a high number of DSBs (Figure 3E). Over time, DSBs were repaired (Figure 3E), satisfying the checkpoint, and by 10 h post-release cells were entering mitosis (Figure 3D), as evidenced by phospho-histone H3 (Hendzel et al., 1997). However, when cells were released from the thymidine-aphidicolin block into CK1δ and CK1ε inhibitors SR-3029 and SR-1227 (Bibian et al., 2013), DSBs persisted for more than 10 h (Figure 3F), and cells failed to enter mitosis (Figure 3G). This is similar to the failure to repair DSBs (Figure 3A-B) and DNA damage checkpoint activation (Figure 2B-C) we observed in *hhp1-as hhp2-as*.

### Hhp1 and Hhp2 phosphorylate multiple substrates to promote HR and NHEJ

To understand how CK1 activity facilitates DSB repair, we focused on the function of Hhp1 and Hhp2 because the CK1-mediated signaling network is simpler in *S. pombe*, and many genetic and biochemical tools are well-established in this model organism. First, we sought to quantify the efficiencies of HR and NHEJ, the two major DSB repair pathways, in the presence and absence of Hhp1 and Hhp2 activity. We used the RDUX200(+) reporter to measure rates of spontaneous HR (Takeda et al., 2008). This reporter gene consists of the *ura4* sequence interrupted by a kanamycin-resistance gene that is flanked by 200-base pair *ura4* repeats. Homologous copies of these *ura4* sequences can recombine to delete the *kanMX6* gene, yielding an intact *ura4^+^*. Colony growth on media lacking uracil is then used to quantify the rate of recombination. Deletion of *rad52*, which is required for HR (Doe et al., 2004; van den Bosch M et al., 2001), prevents nearly all recombination, while inhibiting *hhp1-as hhp2- as* cells significantly reduced the frequency of HR to about half that of uninhibited *hhp1- as hhp2-as* cells and wildtype cells (Figure 4A).

**Figure 4:**
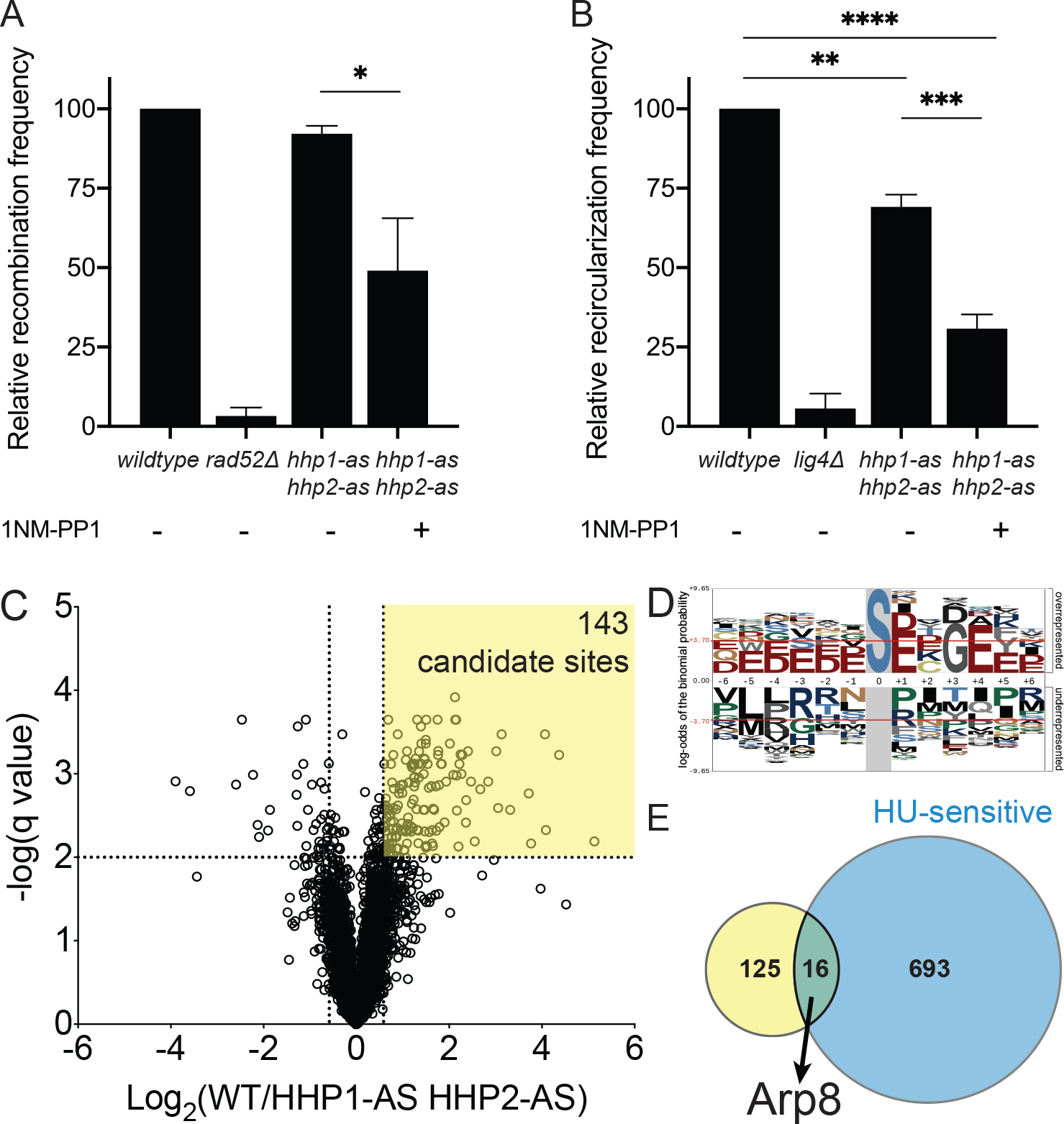
Hhp1 and Hhp2 phosphorylate multiple substrates to promote DSB repair through HR and NHEJ. (A) Analysis of spontaneous HR function using the RDUX200(+) reporter. Data presented as recombination frequency (ratio of colonies grown on selective media to rich media), normalized to *wildtype*. (B) Analysis of NHEJ function using a plasmid recircularization assay. Data presented as recircularization efficiency (ratio of colonies grown from cells transformed with linearized DNA to circular DNA on selective media), normalized to *wildtype*. *, p < 0.05; **, p < 0.01; ***, p < 0.001; ****, p < 0.0001 by one-way ANOVA. (C) Volcano plot comparing statistically significant differences in phosphorylation sites between *hhp1-as hhp1-as* relative to *wildtype* quantified by mass spectrometry-based phosphoproteomics. Dashed lines indicate a Benjamini-Hochberg-corrected q value < 0.01 and a fold change beyond ± 1.5. (D) Linear substrate motif analysis of phosphorylation sites with q < 0.01. (E) Overlap between candidate substrates identified by phosphoproteomics (yellow) and HU-sensitive alleles annotated in PomBase (blue).

To quantify NHEJ function, we used a plasmid recircularization assay, in which a linearized plasmid carrying a *ura4* expression cassette is transformed into cells (Jaendling et al., 2008; Wahls and Davidson, 2008). In order to express *ura4* and survive on uracil-deficient media, cells must recircularize the plasmid using NHEJ. To correct for different transformation efficiencies across strains, cells are also transformed with the circular plasmid, and the linear to circular ratio is calculated for each to yield the frequency of NHEJ. Deletion of *lig4*, the ligase gene required for the final step of NHEJ (Manolis et al., 2001), prevents recircularization (Figure 4B). *hhp1-as hhp2-as* cells, which are slightly hypomorphic even without inhibitor, moderately reduce NHEJ frequency, and treating these cells with 1NM-PP1 further reduces NHEJ to about 30% of wildtype levels (Figure 4B).

These data indicate that both major DSB repair pathways are less efficient in *hhp1* and *hhp2* mutants. Such a limitation to these cells’ repair function is consistent with their poor growth and broad sensitivity to many forms of DNA damage.

Interestingly, Hhp1 and Hhp2 localize diffusely across the nucleus and cytoplasm with a concentration at spindle pole bodies (Elmore et al., 2018), but they did not co-localize with Rad52-GFP foci, even when genotoxic stress was induced (Supplemental Figure 3D). This suggests that their repair function may occur distally from sites of damage.

To identify Hhp1 and Hhp2 substrates that might be involved in these DSB repair functions, we used the *hhp1-as hhp2-as* strain and 1NM-PP1 inhibitor in combination with multiplexed quantitative phosphoproteomics (SL-TMT, Navarrete-Perea et al., 2018). To correct for any changes in protein abundance, we also analyzed total proteome samples (Supplemental Table 2). Each strain was analyzed in biological triplicate. We quantified a total of 4336 phosphorylation sites on 1598 proteins (Supplemental Table 3). Phosphopeptides corresponding to 143 phosphorylation sites on 125 proteins decreased significantly by more than 1.5-fold after inhibition of Hhp1 and Hhp2, representing candidate substrates (Figure 4C). These sites were enriched for the CK1 consensus motif (Flotow et al., 1990; Flotow and Roach, 1991), suggesting that they are likely to contain direct substrates (Figure 4D).

Because loss of Hhp1 and Hhp2 activity results in HU sensitivity and failure to repair DSBs (Figure 2A, 3A-B), we reasoned that loss of Hhp1 and Hhp2 substrates would also confer HU sensitivity. Therefore, to specifically focus on substrates involved in DSB repair, we cross-referenced the candidate substrates identified by phosphoproteomics with alleles that have been annotated with an HU-sensitive phenotype in PomBase, the Global Core Biodata Resource for *S. pombe* (Harris et al., 2022) (Supplemental Table 4). This resulted in 20 phosphorylation sites on 16 proteins (Figure 4E), and we focused on Arp8 for further investigation because this protein contained three phosphorylation sites that were strongly downregulated in *hhp1-as hhp2-as* cells and which all conformed to the CK1 motif (Supplemental Table 2).

Additionally, previous literature established a role for Arp8 in DSB repair (Brahma et al., 2018; Gospodinov et al., 2011; Kashiwaba et al., 2010; Lademann et al., 2017; Morrison et al., 2004; Osakabe et al., 2014; van Attikum et al., 2007), its function could explain defects in both HR and NHEJ (Brahma et al., 2018; Morrison et al., 2004; Poli et al., 2017), and it is conserved from yeast to human. It is important to note, however, that Hhp1 and Hhp2 almost certainly have additional substrates that contribute to DNA repair.

### Arp8, a subunit of the INO80 complex, is an Hhp1 and Hhp2 substrate that is important for DSB repair

Arp8 is a conserved subunit of the INO80 complex, an ATP-dependent nucleosome remodeler that regulates access to the genome during DNA replication, transcription, and DNA repair (Poli et al., 2017). Arp8 facilitates INO80 binding to DNA, including at DSBs (Brahma et al., 2018; Gospodinov et al., 2011; Kashiwaba et al., 2010; Morrison et al., 2004; Osakabe et al., 2014; Tosi et al., 2013; van Attikum et al., 2007). Furthermore, Arp8 promotes end resection at DSBs during both NHEJ (van Attikum et al., 2007) and HR (Gospodinov et al., 2011; Lademann et al., 2017), Rad51 filament formation and eviction of H2AZ during HR (Lademann et al., 2017), and the nucleosome sliding activity of INO80 (Brahma et al., 2018).

To confirm that Arp8 could be phosphorylated by CK1, we tagged *arp8* at the endogenous locus with 3xFLAG, immunoprecipitated the protein from *S. pombe* cells, and incubated with recombinant CK1δ. In accord with our phosphoproteomics results that suggested CK1-mediated phosphorylation of Arp8 in vivo, this in vitro kinase assay also demonstrated phosphorylation of Arp8 (Figure 5A). Next we tested whether the three phosphorylation sites identified by phosphoproteomics (S62, S75, and S87) were indeed where Hhp1 and Hhp2 were phosphorylating Arp8. We expressed and purified recombinant Arp8-WT and Arp8-3A, which has S62, S75, and S87 mutated to alanine, and performed in vitro kinase assays using recombinant Hhp1ΔC and Hhp2ΔC, which have the C-termini truncated to generate highly active kinases (Cegielska et al., 1998; Gietzen and Virshup, 1999; Graves and Roach, 1995; Longenecker et al., 1998).

**Figure 5:**
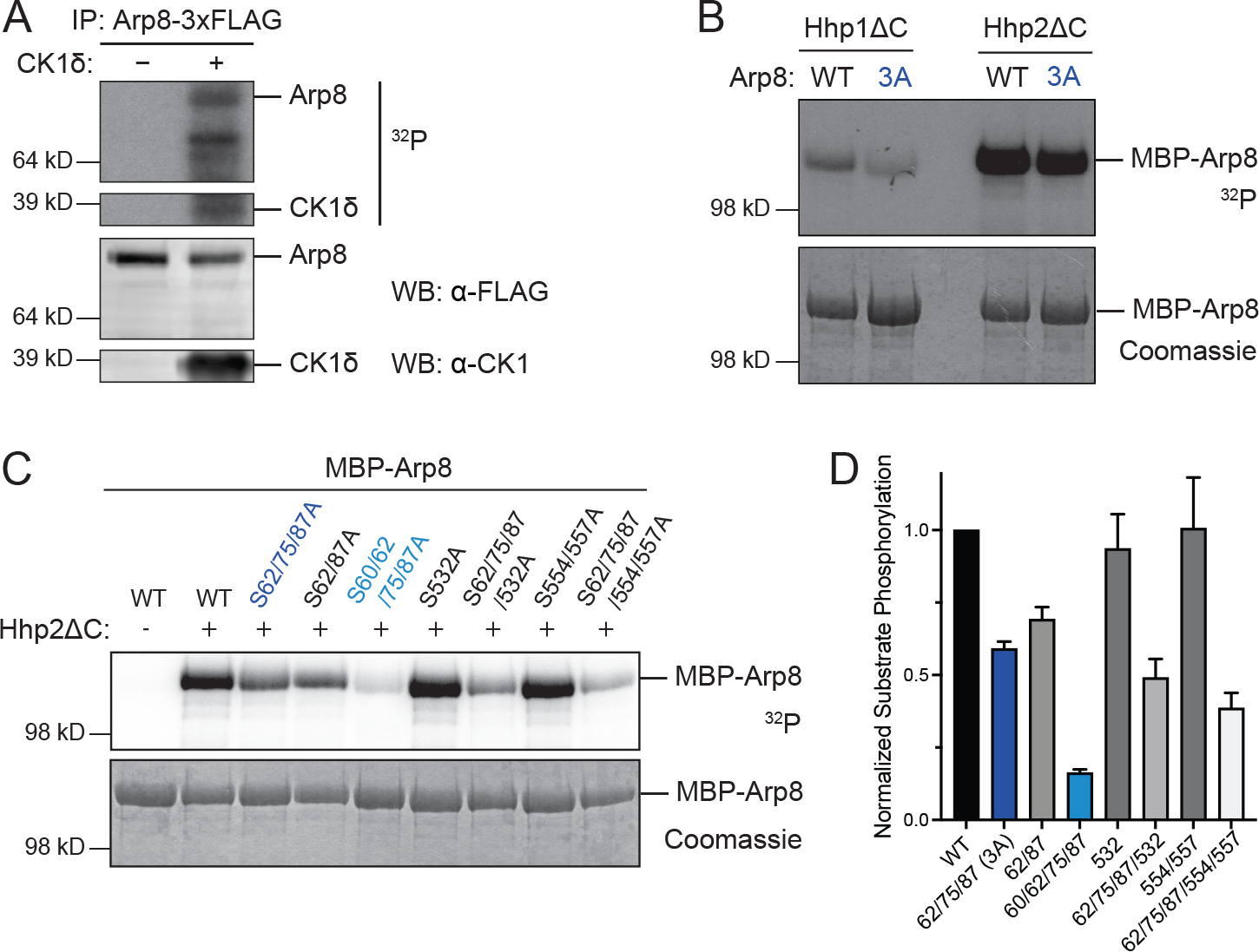
Arp8, a subunit of the INO80 complex, is a substrate of Hhp1 and Hhp2. (A) Arp8-3xFLAG was immunoprecipitated from *S. pombe* then incubated with or without recombinant CK1δ at 30°C for 1 h. Phosphorylated proteins were detected by autoradiography (^32^P) and protein levels by western blot with the indicated antibodies. (B) Recombinant Arp8-WT or Arp8-3A was incubated with recombinant Hhp1ΔC or Hhp2ΔC at 30°C for 1 h. Phosphorylated proteins were detected by autoradiography (^32^P) and protein levels by Coomassie. (C-D) Recombinant Arp8 mutants were incubated with Hhp2ΔC at 30°C for 1 h. Phosphorylated proteins were detected by autoradiography (^32^P) and protein levels by Coomassie. (D) Bars represent means ± SD from 2 independent replicates. ^32^P signal was normalized to protein level for each mutant.

Mutating S62, S75, and S87 to alanine decreased Arp8 phosphorylation by both kinases, indicating that Hhp1 and Hhp2 phosphorylate these sites in vitro (Figure 5B).

Notably, Arp8-3A did not abrogate all phosphorylation, suggesting the presence of additional sites that were not detected in the phosphoproteomics screen.

To identify these remaining sites, we analyzed in vitro phosphorylated Arp8-WT and Arp8-3A by LC-MS^3^. This identified nine phosphorylated serines, including S62 and S87, of which four had high spectral counts and/or the CK1 motif (S60, S532, S554, and S557). We expressed and purified recombinant proteins with these residues mutated to alanine, alone or in combination with the 3A mutations. Because S75 was not detected in the LC-MS^3^ experiment, we also included Arp8-S62A, S87A to compare to Arp8-3A. We incubated the Arp8 phosphorylation site mutants with Hhp2ΔC in vitro and measured the extent of phosphorylation of each substrate (Figure 5C). Arp8-3A reduced phosphorylation to ∼60% of the wildtype level and introducing S60A further reduced phosphorylation to ∼10% (Figure 5D). S60, along with S62 and S87, was also detected in previous phosphoproteomics experiments (Kettenbach et al., 2015; Tay et al., 2019). These data strongly suggest that S60 is an additional Hhp1 and Hhp2 phosphorylation site, and that S60, S62, S75, and S87 are the primary sites on Arp8 that are targeted by Hhp1 and Hhp2. Mutating S532, S554, and S557 slightly decreased overall phosphorylation when combined with 3A, but alone they did not contribute significantly to Arp8 phosphorylation (Figure 5D). These sites also lack the canonical CK1 motif, while S60 has the motif. The contribution of S75 was minimal in vitro, but the in vivo evidence was convincing (Supplemental Table 2), so we reason that S75 may be phosphorylated by Hhp1 and Hhp2 in the cellular context, perhaps following priming at S72.

To confirm that fission yeast Arp8 promotes DNA repair, we grew *arp8Δ* cells on media containing HU and MMS (Supplemental Figure 4A). Consistent with previous literature, *arp8Δ* was sensitive to HU and MMS, as well as to high temperature (Supplemental Figure 4B) (Hogan et al., 2010; Tapia-Alveal et al., 2014). There were no apparent genetic interactions with *hhp1Δ* and *hhp2Δ* (Supplemental Figure 4A-B).

Furthermore, *arp8Δ* had a statistically significant increase in the number of cells with Rad52-GFP foci (Supplemental Figure 4C-D). ∼15% of *arp8Δ* cells had one or more foci, suggesting that loss of *arp8* prevents cells from efficiently repairing DNA damage. The phenotypes of *arp8Δ* are similar to those of *hhp1Δ* and *hhp1Δ hhp2Δ*, suggesting that Arp8 is a substrate through which Hhp1 and Hhp2 may promote DSB repair.

## Discussion

In this study, we have expanded our understanding of Hhp1 and Hhp2’s function in the fission yeast DDR and the consequences of abrogating their activity. Our data demonstrate that Hhp1 and Hhp2 mutants inefficiently repair DSBs, both with and without external sources of genotoxic stress. Cells lacking Hhp1 and Hhp2 activity are still capable of initiating the DDR, including detecting lesions and activating the Rad3- dependent cell cycle checkpoint. Recruitment of early repair proteins such as Rad52 is also intact. Cells deficient in Hhp1 and Hhp2 activity are unable to complete repair, however, and accumulate more and more damage over time, leading to decreased cell survival. The molecular players are conserved from yeast to human, and we see similar accumulation of DSBs in repair foci and cell cycle checkpoint activation in human cells treated with CK1δ and CK1ε inhibitors.

The repair defect in Hhp1 and Hhp2 mutants is pleiotropic, affecting at least two major repair pathways, HR and NHEJ. HR and NHEJ are co-regulated across the cell-cycle, as they compete for broken DNA substrates, and these pathways have been found to share a handful of early DDR proteins, e.g. Mre11, histone H2AX, and Rad3 (Ferreira and Cooper, 2004; Kass and Jasin, 2010; Shrivastav et al., 2008). However, the core proteins responsible for directly engaging repair of DSBs in each pathway are distinct, suggesting that Hhp1 and Hhp2 may have different substrates in HR and NHEJ. This is consistent with the sensitivity of cells lacking Hhp1 and Hhp2 activity to a wide variety of DNA damaging agents. In addition, our proteomics data identified many candidate substrates that are known to have roles in DNA repair (Figure 4E).

We focused on Arp8 because of its well-established role in promoting the function of INO80 during DSB repair by both NHEJ and HR pathways, and because the change in Arp8 phosphorylation upon inhibition of Hhp1 and Hhp2 was pronounced.

The four serines phosphorylated by Hhp1 and Hhp2 all reside in the N-terminus of Arp8. The AlphaFold2 prediction of Arp8 structure (Jumper et al., 2021; Varadi et al., 2021) shows that the N-terminus is poorly predicted and likely flexible; this could be due to or enhanced by different patterns of phosphorylation. In budding yeast, the Arp8 N-terminus was shown to bind to linker DNA between nucleosomes and regulate the nucleosome sliding function of INO80 (Brahma et al 2018). Deleting the N-terminus of Arp8 resulted in HU-sensitivity similar to *arp8Δ* (Brahma et al 2018), demonstrating its essential role in DNA repair. In *S. pombe*, phosphorylation of this region of Arp8 could affect the assembly of the INO80 complex, its localization to DSBs, and/or its nucleosome remodeling activity. We predict that following DNA damage, Hhp1 and Hhp2 phosphorylate Arp8, which opens the chromatin surrounding DSBs through the nucleosome sliding and evicting activity of INO80, making breaks accessible to the repair machinery for HR and NHEJ.

Future studies will determine the mechanism by which Hhp1 and Hhp2 phosphorylation affects the function of Arp8 and INO80, and how this is regulated under different cellular conditions. Future work will also investigate additional substrates that CK1 targets in response to DNA damage. Given the conservation of DNA repair processes and CK1 function from yeast to human, and the similar phenotypes we observed between *S. pombe* and cultured human cells (Figure 3), these findings are likely to also apply to higher eukaryotes.

## Materials and Methods

### Yeast Methods

*S. pombe* strains used in this study (Supplemental Table 1) were grown in yeast extract with supplements (YE) at 32°C unless otherwise indicated. 3’ ORF tagging and gene knockout or disruption was achieved by lithium acetate transformation of PCR-amplified fragments containing regions of homology to the locus of interest (Keeney and Boeke, 1994). Genes were tagged at their 3’ end using pFA6 cassettes as previously described (Bähler et al., 1998). Strains were constructed using standard *S. pombe* crossing and tetrad dissection techniques (Moreno et al., 1991). The ATP analog-sensitive mutants *hhp1-M84G:kanMX6* and *hhp2-M85G:natMX6* were generated in a pIRT2 construct using site-directed mutagenesis, and subsequently integrated over the null using a lithium-acetate protocol. *hhp1-M84G:hphMX6 hhp2-M85G:natMX6* was constructed by swapping the *kanMX6* marker for *hphMX6* by lithium acetate transformation. Inhibition of analog-sensitive Hhp1 and Hhp2 was achieved by exposing cells to 25 μM 1NM-PP1 for 1 h in liquid YE or by growing 2-4 d on YE agar containing 25 μM 1NM-PP1. For growth assays, three 10-fold serial dilutions were made in sterile water starting at 6x10^6^ cells/mL. 3μl of each dilution was spotted on YE plates with the indicated concentrations of DNA damaging agents, and cells were grown at the indicated temperatures.

### Microscopy Methods

Live-cell images of *S. pombe* used for length measurements were acquired with a spinning disk confocal microscope (Ultraview LCI; PerkinElmer, Waltham, MA) equipped with a 63x 1.46 NA PlanApochromat oil immersion objective (Zeiss), an EM- CCD ImagEM X2 camera (Hamamatsu) and μManager software. Strains in the *hhp1- M84G hhp2-M85G* background were treated for 5 h with 25 μM 1NM-PP1 in YE; samples were taken before and after treatment for imaging. Length was measured from a single medial slice in Fiji (Schindelin et al., 2012) by drawing line segments from each pole of the cell to the midpoint of the septum to determine the total cell length at septation.

Representative live-cell images and fixed-cell fluorescence images of *S. pombe* and HeLa cells were acquired using a Personal DeltaVision (Leica Microsystems) that includes a microscope (IX71; Olympus), 60x NA 1.42 Plan Apochromat objective, fixed and live-cell filter wheels, a camera (CoolSNAP HQ2; Photometrics), and softWoRx imaging software (Leica Microsystems). For fixed-cell fluorescence imaging of Rad52-

GFP foci in the *hhp1-M84G hhp2-M85G* background, strains were treated with 12 mM HU for 3 h and then washed into YE containing either 25 μM 1NM-PP1 or DMSO at 32°C. Samples were taken every 30 min for 5 h and fixed with ice-cold 70% ethanol on ice for 15 min. Fixed cells were washed three times in phosphate-buffered saline (PBS) prior to imaging. Eleven z-sections spaced at 0.5 μm were acquired for each image.

Live-cell images of Rad52-GFP foci were acquired using a Zeiss Axio Observer inverted epifluorescence which includes an AxioCam 503 mono camera with Zeiss Plan Apochromat 63x oil (1.46 NA) objective and captured and deconvolved using Zeiss ZEN 3.0 (Blue edition) software. The acquisition setting for Rad52-GFP was set at 100% (intensity) and 150 ms (exposure time) with a z-stack step size of 0.25 µm and a total of 19 Z-slices. Rad52-GFP foci were quantified as described previously (Sabatinos et al., 2012) in Fiji (Schindelin et al., 2012) by applying a Fire heatmap to sum projections of non-deconvolved images. The percentage of cells containing at least one yellow/orange/white focus was counted for each cell population. > 600 cells for each strain were imaged over 2 to 4 independent replicates and pooled. Representative images are deconvolved max projections.

### Human cell culture and synchronization

HeLa cells were cultured in Dulbeco’s modified eagle medium supplemented with 10% fetal bovine serum and 1% penicillin/streptomycin. Cells were synchronized using a sequential thymidine (2.5 mM; Sigma-Aldrich) and aphidicolin (5 μg/mL; Tocris Bioscience) block-and-release protocol. Stock solutions of SR-1227 and SR-3029 were prepared in DMSO; a corresponding volume of DMSO was used for all negative controls.

For microscopy, cells were seeded onto 25 mm coverslips that were contained in six-well plates. Cells were washed once with cold PBS and once with 100% cold methanol, followed by fixation with 100% cold methanol for 15 min at -20°C. Cells were then washed 3x with PBS + 0.1% Tween-20 at RT, followed by blocking with 2% normal goat serum in 0.1% Triton-X100 in PBS for 10 min. Cells were incubated with rabbit anti-γH2AX (Cell Signaling Technology, 2577S), then DRAQ5 (Thermo Fisher Scientific) or DAPI and secondary antibodies (Alexa Fluor Goat-anti Rabbit IgG (H+L), ThermoFisher; 1:500) for 45 minutes at RT. Coverslips were mounted on slides using Prolong Gold antifade mounting media.

For western blotting, ∼40 μg whole cell lysate was incubated with antibodies against γ-tubulin (Sigma-Aldrich, GTU88), phospho-Chk1 (Cell Signaling Technology, 2348S), or phospho-Histone H3 (Sigma, H0412).

### Non-homologous end joining (NHEJ) plasmid recircularization assay

pFY20 (Wahls and Davidson, 2008) was linearized by PstI (New England Biolabs) digest, and complete digestions was confirmed by agarose gel electrophoresis. Prior to transformation, *hhp1-M84G hhp2-M85G* cells were treated with either 25 μM 1NM-PP1 for 1 h at 32°C or a DMSO vehicle control; all other strains were treated with DMSO for 1 h. Cells were transformed with linear, PstI-digested pFY20 or circular, uncut pFY20 via electroporation (Bio-Rad Gene Pulser). Transformants were subsequently grown on Edinburgh minimal media (EMM) lacking uracil at 32°C for 4 d.

Colonies were counted, and the linear/circular ratio (L/C) for each strain was calculated to represent the recircularization efficiency.

### Homologous recombination (HR) assay

The RDUX200(+) reporter system, consisting of an interruption of the endogenous *ura4* locus with a *kanMX6* insertion flanked by 200 bp tandem repeats, was used to measure rates of spontaneous recombination within *hhp1* and *hhp2* mutants (Takeda et al., 2008). The RDUX200(+) reporter was crossed into a *rad52::natMX6* background as well as the ATP analog-sensitive *hhp1-M84G:hphMX6 hhp2-M85G:natMX6* background. Strains were grown to log phase overnight in YE with G418 (100 μg/mL) to eliminate pre-existing recombinants. *hhp1-M84G hhp2-M85G* cells were either treated with 25 μM 1NM-PP1 for 1 h at 32°C or a DMSO vehicle control; all other strains were treated with DMSO for 1 h. Cells were washed with sterile water, diluted, and plated on EMM lacking uracil and YE. Recombination frequency was determined by the ratio of colonies grown on uracil-free media to those grown on YE.

### Protein purification

*Escherichia coli* Rosetta2(DE3)pLysS cells were grown in terrific broth (TB) to an OD of 1.2. Protein production was induced by addition of 0.1 mM IPTG overnight at 17°C. Cells were lysed in lysis buffer (20 mM Tris pH 7.4, 150 mM NaCl, 1 mM EDTA, 0.1% NP-40, 1 mM DTT, 1 mM PMSF, 1.3 mM benzamidine, protease inhibitor tablets [Roche]) using 300 μg/mL lysozyme for 20 minutes followed by sonication. MBP-Arp8 proteins were purified on an MBPTrap HP column (Cytiva) in column buffer (20 mM Tris pH 7.4, 150 mM NaCl, 1 mM EDTA, 1 mM DTT, 1 mM PMSF, 1.3 mM benzamidine) and eluted with maltose (20 mM Tris pH 7.4, 150 mM NaCl, 1 mM EDTA, 1 mM DTT, 1 mM PMSF, 1.3 mM benzamidine, 10 mM maltose, 10% glycerol). Hhp1ΔC and Hhp2ΔC were purified as described previously (Cullati et al., 2022).

### Immunoprecipitation

*S. pombe* strains were grown in YE to log phase. Cell pellets were washed once in NP-40 buffer (10 mM NaPO4 pH = 7.0, 1% NP-40, 150 mM NaCl, 2mM EDTA, 50 mM NaF, 4 μg/mL leupeptin, 100 mM Na3VO4, 1 mM PMSF, 1.3 mM benzamidine, protease inhibitor tablets [Roche], phosphatase inhibitor tablets [Roche]) and lysed by bead disruption using a FastPrep cell homogenizer (MP Biomedicals). The homogenized cell/bead mixture was treated with 500 μL SDS lysis buffer (10 mM sodium phosphate, pH 7.0, 1% SDS, 1 mM DTT, 1 mM EDTA, 50 mM NaF, 100 μM sodium orthovanadate, 1 mM PMSF, 4 μg/mL leupeptin) and incubated at 95°C for 2 min. Lysate was extracted with 800 μL NP-40 buffer and cleared by centrifugation at 17,000 xg for 15 min at 4°C. Arp8-3xFLAG was immunoprecipitated using 2 μg FLAG- M2 (Sigma) at 4°C for 1 h, followed by the addition of Protein A Sepharose beads (GE Healthcare) for 30 min. Proteins were eluted by boiling in SDS-PAGE sample buffer.

Samples were separated by SDS-PAGE, transferred to Immobilon-P polyvinylidene fluoride membrane (Millipore) at 30 V for 1 h, and immunoblotted with mouse anti-FLAG (Sigma) and goat anti-mouse fluorescent antibody (Li-Cor Biosciences), and imaged on an Odyssey CLx (Li-Cor Biosciences).

### In vitro kinase assays

For in vitro kinase assays on immunoprecipitated proteins, beads were equilibrated in 1x PMP buffer (50 mM HEPES, 100 mM NaCl, 2 mM DTT, 0.01% Brij 35, pH 7.5; New England Biolabs). Kinase reactions were performed with 1 μL commercial CK1δ (New England Biolabs) in 1x PMP buffer supplemented with 100 μM unlabeled ATP, 0.5 μCi γ-[^32^P]-ATP, and 10 mM MgCl2 at 30°C for 1 h. For negative control, an equal volume of water instead of CK1δ was added. Kinase reactions were quenched by boiling in SDS-PAGE buffer, separated by SDS-PAGE, and transferred to Immobilon-P polyvinylidene fluoride membrane (Millipore). Phosphoproteins were detected by autoradiography.

200 ng MBP-Hhp1ΔC or MBP-Hhp2ΔC was added to 6 μg MBP-Arp8 and MBP- Arp8-3A in 20 μL 1x PMP buffer supplemented with 10 mM MgCl2, 100 µM unlabeled ATP and 1 µCi γ-[^32^P]-ATP at 30°C for 1 h. Reactions were quenched by boiling in SDS-PAGE buffer, separated by SDS-PAGE, stained with Coomassie, and dried.

Phosphoproteins were detected by autoradiography. For quantification of phosphorylation of Arp8 mutants, ^32^P was quantified on a Typhoon FLA 7000 phosphorimager.

### Mass spectrometry

TCA-precipitated MBP-Arp8 and MBP-Arp8-3A proteins from in vitro kinase assays were subjected to mass spectrometric analysis on an LTQ Velos (Thermo) by 3- phase multidimensional protein identification technology (MudPIT) as previously described (Chen et al., 2013) with the following modifications. Proteins were resuspended in 8M urea buffer (8M urea in 100 mM Tris pH 8.5), reduced with Tris (2- carboxyethyl) phosphine, alkylated with 2-chloro acetamide, and digested with trypsin, chymotrypsin, or elastase. The resulting peptides were desalted by C-18 spin column (Pierce). For the kinase assay samples 6 salt elution steps were used (i.e., 25, 50, 100, 600, 1000, and 5000 mM ammonium acetate) instead of the full 12 steps for TAP samples. Raw mass spectrometry data were first converted to MzXML files using ProteoWizard msconvert (https://proteowizard.sourceforge.io) and then converted to dta files using MzXML2Search (http://tools.proteomecenter.org/wiki/index.php?title= Software:MzXML2Search) before being searched by SEQUEST algorithm (Thermo Fisher Scientific, San Jose, CA, USA; version 27, rev. 12). Scaffold (version 4.8.4) and Scaffold PTM (version 3.2.0) (both from Proteome Software, Portland, OR) were used for data assembly and filtering. The following filtering criteria were used to analyze the phosphorylation sites: minimum of 95% peptide identification probability, minimum of 99% protein identification probability, and minimum of two unique peptides.

### Quantitative phosphoproteomics

Sample processing was performed following the SL-TMT protocol (Navarrete-Perea et al., 2018). Briefly, cell pellets were lysed in 8 M urea complemented with protease and phosphatase inhibitors by bead-beating. After lysis, the protein extracts were centrifuged and the supernatant was obtained. Samples were reduced using 5 mM TCEP for 30 min, alkylated with 10 mM iodoacetamide for 30 min, and the excess of iodoacetamide was quenched using DTT. After protein quantification, 200 µg of protein were chloroform-methanol precipitated and reconstituted in 200 µL of 200 mM EPPS (pH 8.5). Protein was digested using Lys-C overnight at room temperature followed by trypsin for 6 h at 37°C, both at a 100:1 protein-to-protease ratio. After digestion, the samples were labeled using the TMTpro16 reagents for 60 min (Thermo), the reactions were quenched using hydroxylamine (final concentration of 0.3% v/v) for 20 min. After label check, the samples were combined equally and desalted. Phosphopeptides were enriched using the Pierce High-Select Fe-NTA Phosphopeptide Enrichment kit following manufacturer’s instructions. The phosphopeptides were eluted in a tube containing 100 µL of 10% formic acid and dried in a vacuum centrifuge. The unbound fraction was retained and fractionated using basic pH reversed-phase (BPRP) HPLC, the resulting 96 fractions were consolidated into 24, and 12 were processed in the mass spectrometer. The phosphopeptides were fractionated using the Pierce High pH Reversed-Phase Peptide Fractionation Kit following manufacturer’s instructions. The phosphopeptides were eluted using the following ACN concentrations: 7.5, 10, 12.5, 15, 17.5, 20, 22.5, 25, 27.5, 30, 40 and 50% ACN, then the samples were combined intro 3 final fractions: a) 7.5, 20, 22.5 and 50, b) 10, 17.5, 25 and 40, c) 12.5, 15, 27.5 and 30. All data were collected on an Orbitrap Eclipse mass spectrometer coupled to a Proxeon NanoLC-1000 UHPLC. The peptides were separated using a 100 μm capillary column packed with ≈35 cm of Accucore 150 resin (2.6 μm, 150 Å; ThermoFisher Scientific). For BPRP fractions, the data were collected using a DDA-SPS-MS3 method with online real-time database searching (RTS) coupled with FAIMS (Schweppe et al., 2020a, 2020b). Each fraction was eluted using a 90 min method over a gradient from 6% to 30% ACN. Peptides were ionized with a spray voltage of 3,000 kV. The instrument method included Orbitrap MS1 scans (resolution of 120,000; mass range 400−1400 m/z; automatic gain control (AGC) target 2x10^5^, max injection time of 50 ms and ion trap MS2 scans (CID collision energy of 35%; AGC target 1x10^4^; rapid scan mode; max injection time of 120 ms). RTS was enabled and quantitative SPS-MS3 scans (resolution of 50,000; AGC target 2.5x10^5^; max injection time of 250 ms) were processed through Orbiter with a real-time false discovery rate filter implementing a modified linear discriminant analysis.

Phosphopeptides were analyzed using FAIMS/hrMS2 following our optimized workflow for multiplexed phosphorylation analysis (Schweppe et al., 2020a, 2020b). Briefly, the Thermo FAIMS Pro device was operated with default parameters. No additional gas was used for desolvation. The DV circuitry was auto-tuned, which independently tunes each of the sine waves and phase shifts one of the waveforms by π/2 to assemble a bisinusoidal waveform with a high amplitude of −5000 V at a 3 MHz frequency. The three phosphopeptide fractions were analyzed twice in the mass spectrometer, once with a method incorporating two CVs (CV= -45 and -70V) and the other with three CVs (CV= -40V, -60V and -80V) using a 2 h method using a gradient of 6% to 30% B.

Raw files were first converted to mzXML. Database searching included all *Schizosaccharomyces pombe* entries from UniProt (downloaded December 2020). The database was concatenated with one composed of all protein sequences in the reversed order and a list of common contaminant proteins was also included. Searches were performed using a 50 ppm precursor ion tolerance and 0.9 Da (low-resolution MS2) or 0.03 Da (high-resolution MS2) product ion tolerance (Beausoleil et al., 2006; Elias and Gygi, 2007; Huttlin et al., 2010). TMTpro on lysine residues and peptide N termini (+304.2071 Da for TMTpro) and carbamidomethylation of cysteine residues (+57.0215 Da) were set as static modifications (except when testing for labeling efficiency, when the TMTpro modifications are set to variable). Oxidation of methionine residues (+15.9949 Da) was set as a variable modification. For phosphopeptide analysis, +79.9663 Da was set as a variable modification on serine, threonine, and tyrosine residues. The mass spectrometry proteomics data have been deposited to the ProteomeXchange Consortium via the PRIDE (Perez-Riverol et al., 2019) partner repository with the data set identifier PXD033223.

t tests were used to compare each protein or phosphorylation site measurement, and a Benjamini−Hochberg multiple testing correction was applied. Statistical analysis and volcano plots were generated in Graph Pad Prism v8. Linear substrate motif analysis was performed on phosphopeptides with q < 0.01 using pLogo (O’Shea et al., 2013).

### Statistical analysis

Statistical analysis was performed using GraphPad Prism v8.

## Acknowledgements

We thank Dr. Alaina Willet for critical comments on the manuscript. S.N.C. was supported by the Integrated Biological Systems Training in Oncology Program (T32- CA119925). This work was funded by R35-GM131799 to K.L.G.

## Author contributions

S.N.C. and K.L.G. conceptualized the study. E.Z., Y.S., R.X.G., Z.C.E., and L.R. performed experiments. S.N.C., J.S.C, J.N.P, and S.P.G. prepared and analyzed phosphoproteomics samples. S.N.C., E.Z., Y.S., and K.L.G. wrote the original draft. All authors revised the manuscript.

