## Supplemental Figures 1-4 and Supplemental Table 1 for "Fission yeast CK1 promotes DNA double-strand break repair through both homologous recombination and non-homologous end joining"

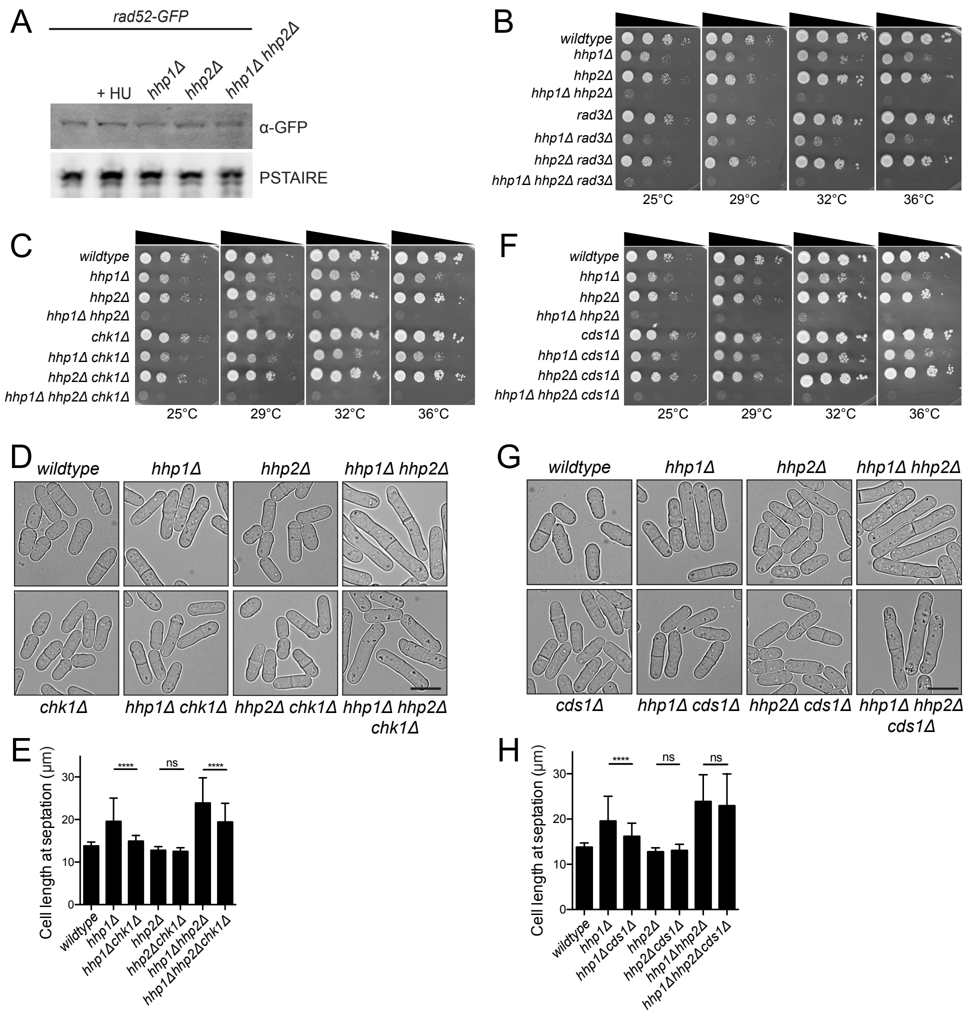


Supplemental Figure 1

**Supplemental Figure 1: Checkpoint kinases suppress the cell length phenotype but do not rescue viability of *hhp1Δ* and *hhp1Δ hhp2Δ* cells.** (A) Anti-GFP western blot of cell lysates from the indicated strains, with anti-Cdc2 (PSTAIRE) as the loading control. (B, C, F) Serial 10-fold dilutions of the indicated strains were spotted on YE and incubated at the indicated temperatures. (D-E, G-H) Live cells were imaged by DIC and length at septation was measured. For *wildtype* and *hhp1Δ hhp2Δ* strains, $n \geq 40$ cells in each of 6 replicates. For all other strains, $n \geq 40$ cells in each of 3 replicates. *wildtype, hhp1Δ, hhp2Δ*, and *hhp1Δ hhp2Δ* data are shared with Figures 1D, S1E, and S1F. Bars in E and H represent means ± SD. ****, $p < 0.0001$ by one-way ANOVA; ns, not significant. Scale bar: 10 μm.

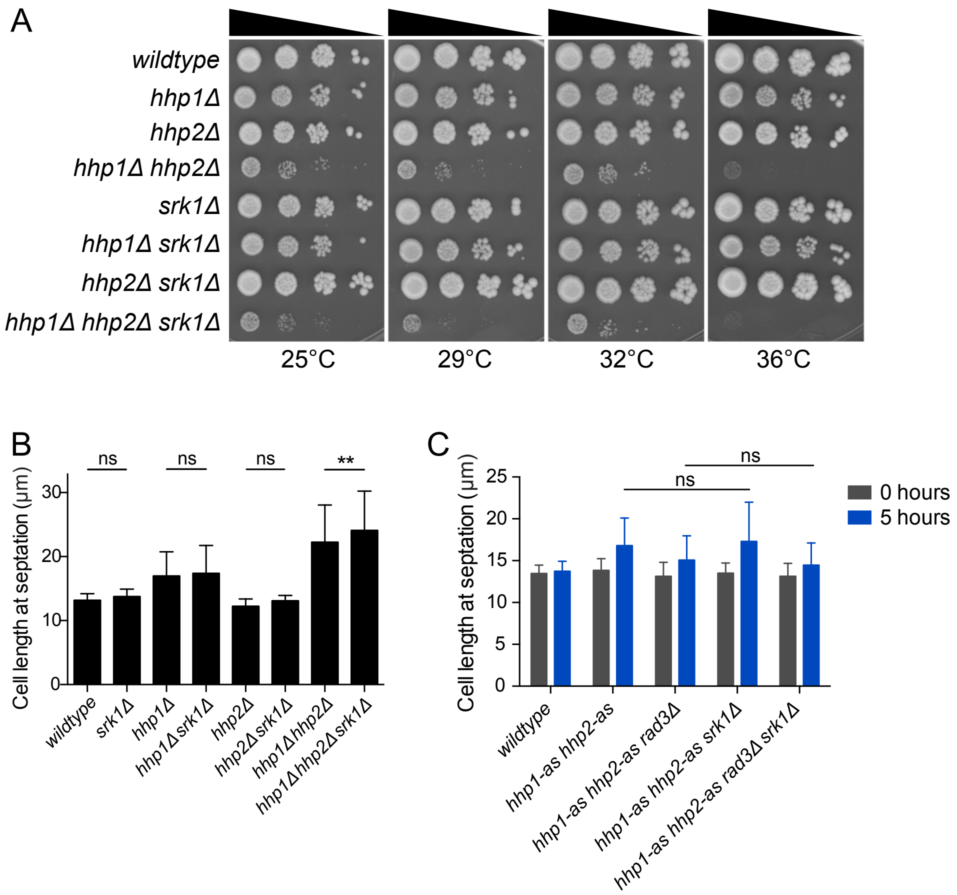


Supplemental Figure 2

**Supplemental Figure 2: The *hhp1Δ* and *hhp1Δ hhp2Δ* length phenotype is not suppressed by deletion of stress-activated checkpoint kinase *srk1*.** (A) Serial 10-fold dilutions of the indicated strains were spotted on YE and incubated at the indicated temperatures. (B) Live cells were imaged by DIC and length at septation was measured, $n \geq 38$ cells in each of 3 replicates. Bars represent means ± SD. **, $p < 0.01$ by one-way ANOVA; ns, not significant. (C) Deletion of *srk1* does not significantly alter cell length. Log-phase cultures from the indicated strains were treated with 25 μM 1NM-PP1 for 5 h. Live cells were imaged by DIC and length at septation was measured at 0 and 5 h, $n \geq 50$ cells in each of 3 replicates. Bars represent means ± SD. p values determined by one-way ANOVA; ns, not significant.

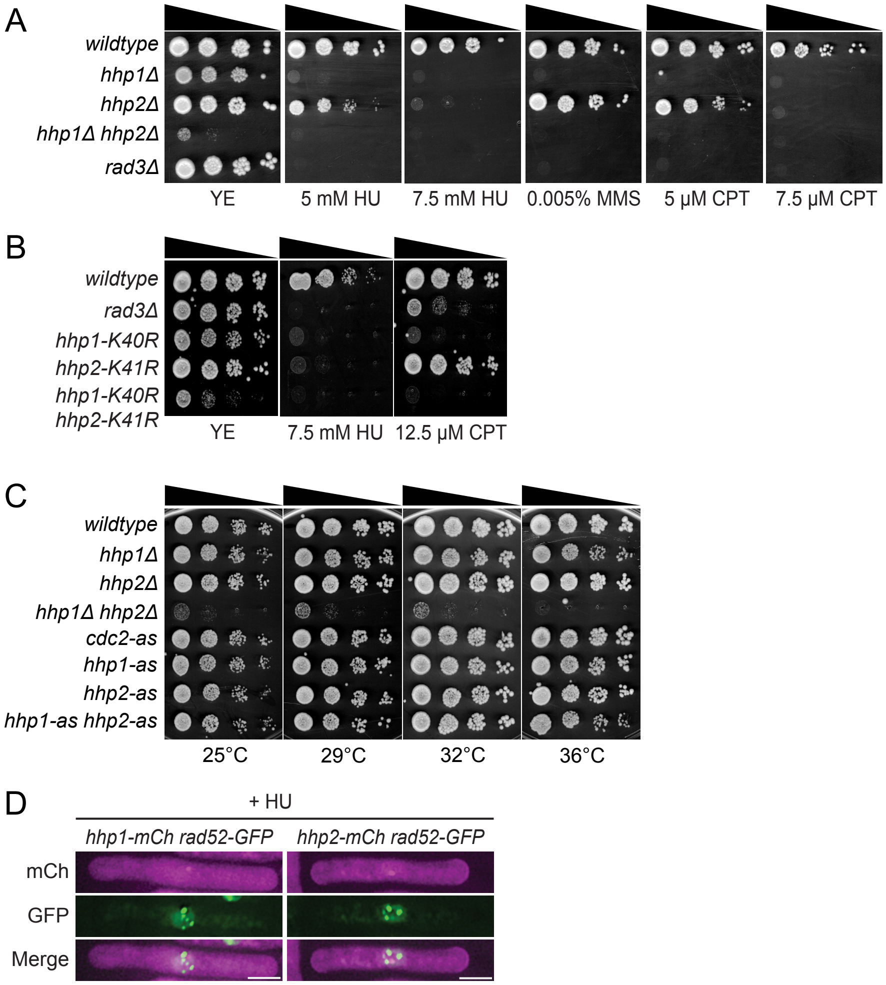


Supplemental Figure 3

**Supplemental Figure 3: Hhp1 and Hhp2 kinase activity is required for cells to survive persistent DNA damage.** (A-C) Serial 10-fold dilutions of the indicated strains were spotted on YE containing drugs that induce DNA damage and incubated at 32°C. *hhp1-K40R* and *hhp2-K41R* kinase-dead mutants (B) were as sensitive as knockouts (A) to replication stress and DSBs. Analogue-sensitive strains (C) grew normally without 1NM-PP1. (D) Fluorescence microscopy of the indicated strains following treatment with hydroxyurea. Hhp1 and Hhp2 do not colocalize with Rad52-GFP at DSBs.

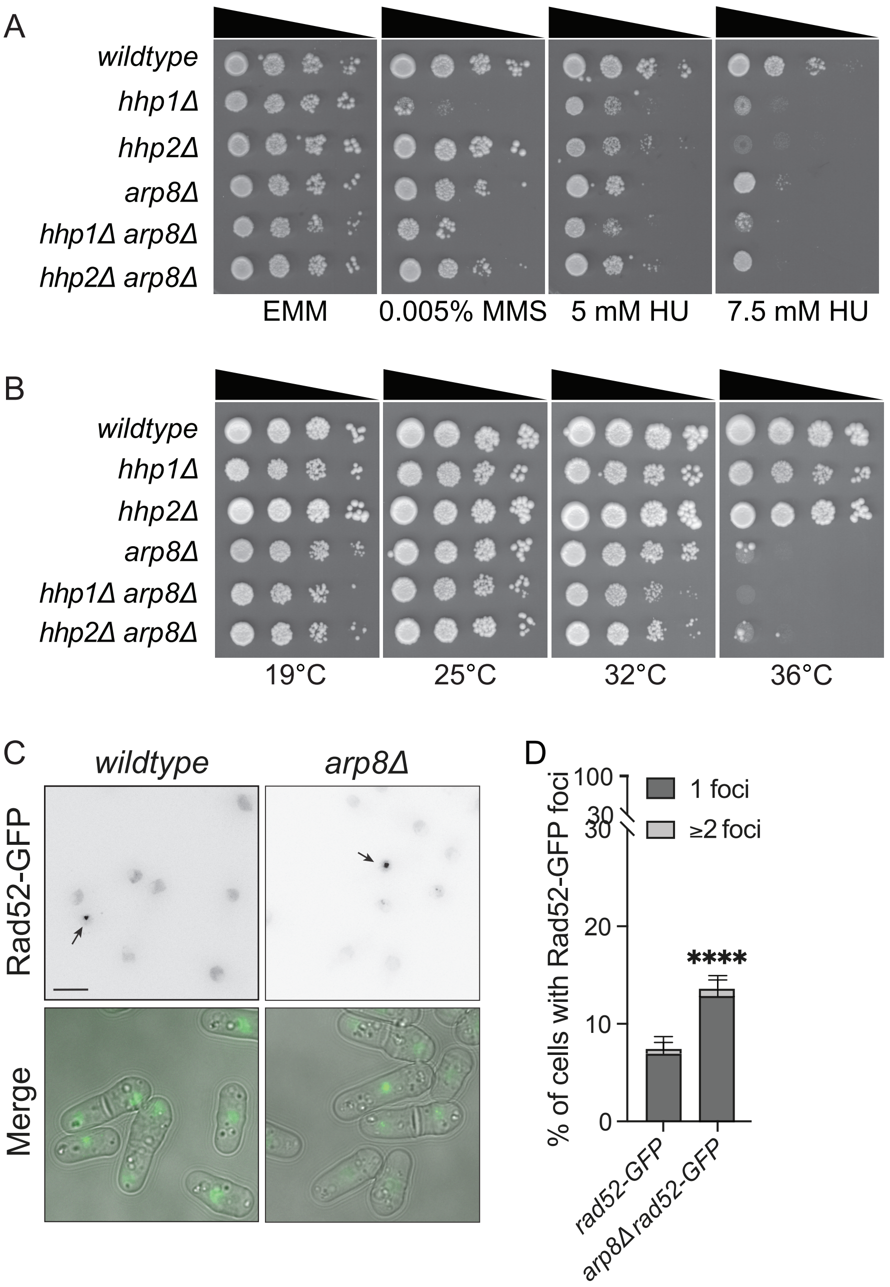


Supplemental Figure 4

**Supplemental Figure 4: Arp8 is important for DSB repair.** (A) Serial 10-fold dilutions of the indicated strains were spotted on EMM containing drugs that induce DNA damage and incubated at 32°C. (B) Serial 10-fold dilutions of the indicated strains were spotted on YE and incubated at the indicated temperatures. (C-D) *arp8Δ* exhibits increased basal levels of Rad52-GFP foci compared to *wildtype*. (C) Fluorescence microscopy and DIC images of the indicated strains. (D) Rad52-GFP foci in > 600 cells for each strain were counted over 2 to 4 biological replicates. Error bars indicate ± 95% CI. ****, $p < 0.0001$ by Chi-square. Wildtype data is shared with Figure 1A-B.

**Supplemental Table 1: *S. pombe* strains used in this study.**

| Strain | Genotype | Source |
|---|---|---|
| Figure 1 | | |
| 16715 | *rad52-GFP:kanMX6 ade6-M210 ura4-D18 leu1-32 h-* | This study |
| 16767 | *rad52-GFP:kanMX6 hhp1Δ::ura4*$^{+}$ *ade6-M210 ura4-D18 leu1-32 h-* | This study |
| 17076 | *rad52-GFP:kanMX6 hhp2Δ::ura4*$^{+}$ *ade6-M210 ura4-D18 leu1-32 h+* | This study |
| 17126 | *rad52-GFP:kanMX6 hhp1Δ::ura4*$^{+}$ *hhp2Δ::ura4*$^{+}$ *ade6-M210 ura4-D18 leu1-32 h?* | This study |
| 83 | *ade6-M210 leu1-32 h-* | Lab stock |
| 17578 | *hhp1Δ::ura4*$^{+}$ *ade6-M210 ura4-D18 leu1-32 h-* | Bimbo et al. 2005 |
| 7683 | *hhp2Δ::ura4*$^{+}$ *ade6-M210 ura4-D18 leu1-32 h+* | Bimbo et al. 2005 |
| 2747-2 | *hhp1Δ::ura4*$^{+}$ *hhp2Δ::kanMX6 ade6-M210 ura4-D18 leu1-32 h-* | Elmore et al. 2018 |
| 2748-2 | *rad3Δ::ura4*$^{+}$ *ade6-M210 ura4-D18 leu1-32 h+* | Bimbo et al. 2005 |
| 17345 | *rad3Δ::kanMX6 hhp1Δ::ura4*$^{+}$ *ade6-M210 ura4-D18 leu1-32 h-* | This study |
| 14478 | *rad3Δ::ura4*$^{+}$ *hhp2Δ::kanMX6 ade6-M210 ura4-D18 leu1-32 h+* | This study |
| 3995-2 | *rad3Δ::ura4*$^{+}$ *hhp1Δ::ura4*$^{+}$ *hhp2Δ::kanMX6 ade6-M210 ura4-D18 leu1-32 h?* | This study |
| Figure 2 | | |
| 246 | *ade6-M210 ura4-D18 leu1-32 h-* | Lab stock |
| 4969 | *cdc2-F84G ade6-M210 ura4-D18 leu1-32 h+* | Lab stock |
| 17707 | *hhp1-M84G:kanMX6 ade6-M210 ura4-D18 leu1-32 h+* | Cullati et al. 2022 |
| 17596 | *hhp2-M85G:natMX6 ade6-M210 ura4-D18 leu1-32 h-* | Cullati et al. 2022 |
| 17712 | *hhp1-M84G:kanMX6 hhp2-M85G:natMX6 ade6-M210 ura4-D18 leu1-32 h-* | This study |
| 4030-2 | *rad3Δ::ura4*$^{+}$ *hhp1-M84G:kanMX6 hhp2-M85G:natMX6 ade6-M210 ura4-D18 leu1-32 h?* | This study |
| 4007-2 | *chk1Δ::ura4*$^{4}$ *hhp1-M84G:kanMX6 hhp2-M85G:natMX6 ade6-M210 ura4-D18 leu1-32 h-* | This study |
| 4036-2 | *cds1Δ::ura4*$^{+}$ *hhp1-M84G:kanMX6 hhp2-M85G:natMX6 ade6-M210 ura4-D18 leu1-32 h?* | This study |

| | | |
|---|---|---|
| Figure 3 | | |
| 18821 | rad52-GFP:kanMX6 hhp1-M84G:kanMX6 hhp2-M85G:natMX6 ade6-M210 ura4-D18 leu1-32 h? | This study |
| Figure 4 | | |
| 19361 | ura4::RDUX200(kanMX6) leu1-32 h- | Manjon et al. 2017 |
| 19667 | rad52Δ::natMX6 ura4::RDUX200(kanMX6) leu1-32 h- | This study |
| 18408 | hhp1-M84G:hphMX6 hhp2-M85G:natMX6 ura4::RDUX200(kanMX6) leu1-32 h? | This study |
| 246 | ade6-M210 ura4-D18 leu1-32 h- | Lab stock |
| 17049 | lig4Δ::kanMX6 ade6-M210 ura4-D18 leu1-32 h+ | This study |
| 17712 | hhp1-M84G:kanMX6 hhp2-M85G:natMX6 ade6-M210 ura4-D18 leu1-32 h- | This study |
| Supplemental Figure 1 | | |
| 83 | ade6-M210 leu1-32 h- | Lab stock |
| 17578 | hhp1Δ::ura4$^+$ ade6-M210 ura4-D18 leu1-32 h- | Bimbo et al. 2005 |
| 7683 | hhp2Δ::ura4$^+$ ade6-M210 ura4-D18 leu1-32 h+ | Bimbo et al. 2005 |
| 2747-2 | hhp1Δ::ura4$^+$ hhp2Δ::kanMX6 ade6-M210 ura4-D18 leu1-32 h- | Elmore et al. 2018 |
| 2748-2 | rad3Δ::ura4$^+$ ade6-M210 ura4-D18 leu1-32 h+ | Bimbo et al. 2005 |
| 17345 | rad3Δ::kanMX6 hhp1Δ::ura4$^+$ ade6-M210 ura4-D18 leu1-32 h- | This study |
| 14478 | rad3Δ::ura4$^+$ hhp2Δ::kanMX6 ade6-M210 ura4-D18 leu1-32 h+ | This study |
| 3995-2 | rad3Δ::ura4$^+$ hhp1Δ::ura4$^+$ hhp2Δ::kanMX6 ade6-M210 ura4-D18 leu1-32 h? | This study |
| 511 | chk1Δ::ura4$^+$ ade6-M216 ura4-D18 leu1-32 h+ | Bimbo et al. 2005 |
| 17346 | chk1Δ::kanMX6 hhp1Δ::ura4$^+$ ade6-M210 ura4-D18 leu1-32 h- | This study |
| 17667 | chk1Δ::kanMX6 hhp2Δ::ura4$^+$ ade6-M210 ura4-D18 leu1-32 h+ | This study |
| 4001-2 | chk1Δ::kanMX6 hhp1Δ::ura4$^+$ hhp2Δ::ura4$^+$ ade6-M210 ura4-D18 leu1-32 h? | This study |
| 2931-2 | cds1Δ::ura4$^+$ ade6-M210 ura4-D18 leu1-32 h- | Bimbo et al. 2005 |
| 17347 | cds1Δ::kanMX6 hhp1Δ::ura4$^+$ ade6-M210 ura4-D18 leu1-32 h+ | This study |

| | | |
|---|---|---|
| 2932-2 | *cds1Δ::kanMX6 hhp2Δ::ura4*$^+$ *ade6-M210 ura4-D18 leu1-32 h-* | This study |
| 3996-2 | *cds1Δ::kanMX6 hhp1Δ::ura4*$^+$ *hhp2Δ::ura4*$^+$ *ade6-M210 ura4-D18 leu1-32 h?* | This study |
| Supplemental Figure 2 | | |
| 83 | *ade6-M210 leu1-32 h-* | Lab stock |
| 17578 | *hhp1Δ::ura4*$^+$ *ade6-M210 ura4-D18 leu1-32 h-* | Bimbo et al. 2005 |
| 7683 | *hhp2Δ::ura4*$^+$ *ade6-M210 ura4-D18 leu1-32 h+* | Bimbo et al. 2005 |
| 2747-2 | *hhp1Δ::ura4*$^+$ *hhp2Δ::kanMX6 ade6-M210 ura4-D18 leu1-32 h-* | Elmore et al. 2018 |
| 4015-2 | *srk1Δ::kanMX6 ade6-M210 ura4-D18 leu1-32 h+* | This study |
| 4032-2 | *srk1Δ::kanMX6 hhp1Δ::ura4*$^+$ *ade6-M210 ura4-D18 leu1-32 h+* | This study |
| 4050-2 | *srk1Δ::kanMX6 hhp2Δ::ura4*$^+$ *ade6-M210 ura4-D18 leu1-32 h-* | This study |
| 4063-2 | *srk1Δ::kanMX6 hhp1Δ::ura4*$^+$ *hhp2Δ::ura4*$^+$ *ade6-M210 ura4-D18 leu1-32 h?* | This study |
| 17712 | *hhp1-M84G:kanMX6 hhp2-M85G:natMX6 ade6-M210 ura4-D18 leu1-32 h-* | This study |
| 4030-2 | *rad3Δ::ura4*$^+$ *hhp1-M84G:kanMX6 hhp2-M85G:natMX6 ade6-M210 ura4-D18 leu1-32 h?* | This study |
| 4072-2 | *srk1Δ::kanMX6 hhp1-M84G:kanMX6 hhp2-M85G:natMX6 ade6-M210 ura4-D18 leu1-32 h+* | This study |
| 4081-2 | *srk1Δ::kanMX6 rad3Δ::ura4*$^+$ *hhp1-M84G:kanMX6 hhp2-M85G:natMX6 ade6-M210 ura4-D18 leu1-32 h?* | This study |
| Supplemental Figure 3 | | |
| 83 | *ade6-M210 leu1-32 h-* | Lab stock |
| 17578 | *hhp1Δ::ura4*$^+$ *ade6-M210 ura4-D18 leu1-32 h-* | Bimbo et al. 2005 |
| 7683 | *hhp2Δ::ura4*$^+$ *ade6-M210 ura4-D18 leu1-32 h+* | Bimbo et al. 2005 |
| 2747-2 | *hhp1Δ::ura4*$^+$ *hhp2Δ::kanMX6 ade6-M210 ura4-D18 leu1-32 h-* | Elmore et al. 2018 |
| 2748-2 | *rad3Δ::ura4*$^+$ *ade6-M210 ura4-D18 leu1-32 h+* | Bimbo et al. 2005 |
| 19337 | *hhp1-K40R:kanMX6 ura4-D18 leu1-32 h-* | Elmore et al. 2018 |
| 19349 | *hhp2-K41R:kanMX6 ade6-M210 ura4-D18 leu1-32 h+* | Elmore et al. 2018 |

| | | |
|---|---|---|
| 19342 | *hhp1-K40R:kanMX6 hhp2-K41R:kanMX6 ade6-M210 ura4-D18 leu1-32 h?* | Elmore et al. 2018 |
| 246 | *ade6-M210 ura4-D18 leu1-32 h-* | Lab stock |
| 4969 | *cdc2-F84G ade6-M210 ura4-D18 leu1-32 h+* | Lab stock |
| 17707 | *hhp1-M84G:kanMX6 ade6-M210 ura4-D18 leu1-32 h+* | Cullati et al. 2022 |
| 17596 | *hhp2-M85G:natMX6 ade6-M210 ura4-D18 leu1-32 h-* | Cullati et al. 2022 |
| 17712 | *hhp1-M84G:kanMX6 hhp2-M85G:natMX6 ade6-M210 ura4-D18 leu1-32 h-* | This study |
| 19150 | *rad52-GFP:kanMX6 hhp1-mCherry:kanMX6 ade6-M210 ura4-D18 leu1-32 h?* | This study |
| 19151 | *rad52-GFP:kanMX6 hhp2-mCherry:kanMX6 ade6-M210 ura4-D18 leu1-32 h?* | This study |
| Supplemental Figure 4 | | |
| 83 | *ade6-M210 leu1-32 h-* | Lab stock |
| 17578 | *hhp1Δ::ura4$^+$ ade6-M210 ura4-D18 leu1-32 h-* | Bimbo et al. 2005 |
| 7683 | *hhp2Δ::ura4$^+$ ade6-M210 ura4-D18 leu1-32 h+* | Bimbo et al. 2005 |
| 7280-2 | *arp8Δ::ura4$^+$ ade6-M210 ura4-D18 leu1-32 h-* | This study |
| 7398-2 | *hhp1Δ::ura4$^+$ arp8Δ::ura4$^+$ ade6-M210 ura4-D18 leu1-32 h+* | This study |
| 7399-2 | *hhp2Δ::ura4$^+$ arp8Δ::ura4$^+$ ade6-M210 ura4-D18 leu1-32 h-* | This study |
| 16715 | *rad52-GFP:kanMX6 ade6-M210 ura4-D18 leu1-32 h-* | This study |
| 7409-2 | *rad52-GFP:kanMX6 arp8Δ::ura4$^+$ ade6-M210 ura4-D18 leu1-32 h-* | This study |

**Supplemental Table 2: Proteins quantified in the proteome dataset.**

**Supplemental Table 3: Phosphorylation sites quantified in the phosphoproteome dataset.**

**Supplemental Table 4: Gene set with HU-sensitive phenotype.**
